# Prevention of Transgene Silencing During Human Pluripotent Stem Cell Differentiation

**DOI:** 10.1101/2025.04.07.647695

**Authors:** Takeshi Uenaka, Alan Bortoncello Napole, Aninda Dibya Saha, Duo Sun, Angelina Singavarapu, Liz Calzada, Jiahui Chen, Lena Erlebach, Amanda McQuade, Daniel Ramos, Alessandra Rigamonti, Lisa Salazar, Avi J. Samelson, Kamilla Sedov, Natalie J. Welsh, Katleen Wild, Qianxin Wu, Ernest Arenas, Andrew R. Bassett, Martin Kampmann, Deborah Kronenberg-Versteeg, Florian T. Merkle, Birgitt Schüle, Leslie M. Thompson, William C. Skarnes, Michael E. Ward, Marius Wernig

## Abstract

While high and stable transgene expression can be achieved in undifferentiated pluripotent stem cells, conventional transgene expression systems are often silenced upon differentiation. Silencing occurs with both randomly integrated transgenes, introduced via transposase or lentiviral methods, and with transgenes targeted to specific genomic sites, including at commonly used safe harbor loci. The challenge to robustly express experimental transgenes in differentiated pluripotent stem cells is a major bottleneck in the field for applications such as CRISPR screening. Here, we conducted a comparative analysis to systematically evaluate the impact of various promoters, transcriptional regulatory elements, insulators, and genomic integration sites on transgene silencing during neuronal differentiation. Our findings reveal that specific combinations of promoters and transcriptional stability elements are able to prevent transgene silencing during differentiation, whereas chromatin insulators had less impact on silencing and three novel safe harbor integration sites performed similarly to the *CLYBL* locus. Guided by these insights we developed the PiggyBac vector TK4, which showed complete resistance to transgene silencing across various neuronal and microglial differentiation protocols from six different pluripotent stem cell lines, as independently confirmed by seven different laboratories. This construct will be highly useful for assays requiring stable transgene expression during differentiation, and holds the potential for broad applications in various research fields.

## INTRODUCTION

Differentiating human embryonic stem (ES) or induced pluripotent stem (iPS) cells into various cell types is an essential technology for studying human development, disease mechanisms, and potential therapeutic applications^1^. Achieving these goals often involves engineering stem cells by overexpressing transgenes or downregulating endogenous genes, which are indispensable techniques. However, a significant challenge for the field is that these transgenes are frequently silenced. This silencing occurs in both undifferentiated stem cells^2^ and cells undergoing differentiation into other cell types including cardiomyocytes^3^, endothelium^4^, myeloid cells^5^, and neurons^6^.

Several papers have reported mechanisms contributing to transgene silencing. First, specific sequences within the vector, such as long terminal repeats (LTRs) in retroviral vectors^7^, promoters^8^, or intron-less transgenes via human silencing hub (HUSH) complex^9^, can recruit DNA modifications such as DNA methylation or histone deacetylation to silence genes^10,11^. Second, integration of the vector into heterochromatin regions can lead to spreading of transcriptional repressive epigenetic modifications^12,13^. Third, posttranscriptional mechanims involving double-stranded RNA have been described in particular in plant and fungal cells^14^. Incorporating insulator elements into the expression vector^2, 15^ or targeting expression cassettes into commonly used safe-harbor loci such as the adeno-associated virus integration site 1 (AAVS1)^16^ or citrate lyase beta-like (CLYBL)^17^ locus allowed more stable gene expression over time in proliferating, undifferentiated pluripotent stem cells. However, epigenetic marks are known to be tissue-specific^18^ and high and stable gene expression in pluripotent cells accomplished by insulators or gene targeting does not translate to sustained expression during lineage differentiation.

The inability to maintain exogenous gene expression throughout differentiation poses a significant obstacle for many experimental approaches, including cell labeling with fluorescent reporters and overexpressing functional proteins like disease mutant proteins^19^, GCaMP^20^, and dead Cas9^21^. Most critically, this silencing issue compromises the robustness of CRISPR/Cas9-related genetic screens, which would otherwise be ideally suited for use with pluripotent stem cell platforms. Therefore, there is an urgent need to overcome transgene silencing that occurs during differentiation.

Here, we present the PiggyBac construct TK4 that can prevent transgene silencing during the differentiation of human pluripotent cells into neurons or microglia. Through systematic comparison, we identify a combination of transcriptional elements that effectively overcomes silencing during differentiation.

## DESIGN

A critical bottleneck in the field is the pronounced silencing of experimental transgenes during human pluripotent stem cell differentiation. To overcome this limitation, we took a two-pronged approach: 1) to test several relevant regulatory DNA elements in expression vectors and 2) to identify an optimal genomic integration site. Both approaches are known to affect gene expression but a systematic evaluation of combinations of regulatory elements and comparing different genomic integration sites has not been performed yet for differentiation of pluripotent stem cells.

## RESULTS

### Establishment of a PiggyBac vector that can prevent transgene silencing during neuronal differentiation

When human pluripotent stem cells are differentiated into neurons, transgene expression is substantially silenced, either when expressed from random integrations or from defined genomic sites such as the *CLYBL* locus (**Figures 1A-1C**). To identify an expression system that may avoid silencing, we empirically designed a multi-element PiggyBac vector (TK4 vector, **Figure 1D**). This vector is highly modular as all elements of the transcriptional expression cassette are flanked by unique restriction enzymes that allow facile element exchange. To quantify the degree of transcriptional silencing we introduced a CAG-EGFP-WPRE-pA expression cassette in one direction. We had previously determined that EGFP can be reliably used to measure silencing (**Figure 1C**). In the opposite direction we introduced an EF1α-puro-T2A-mTagBFP2-pA construct for drug-selection of cells with successful vector integration. Upstream of the EGFP expression cassette we further introduced the long cHS4^15^ and the SRF-UCOE insulator elements^22^. In addition, we kept the two core regions of the cHS4 insulator element that were already contained in the original vector and flanked the entire inserted DNA cassette.

**Figure 1.**
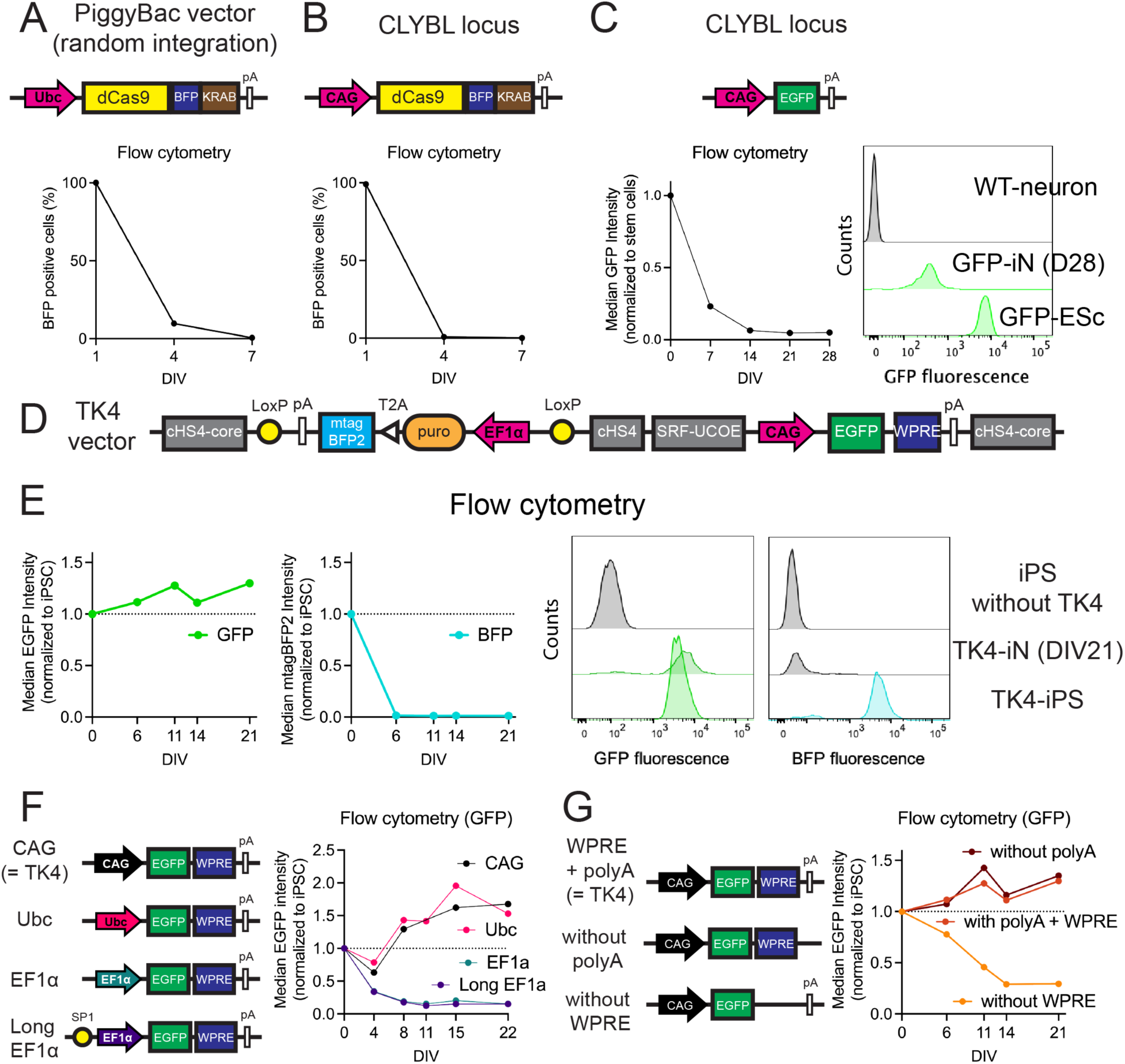
Promoter and WPRE sequences are the key elements in the TK4 PiggyBac vector that prevent silencing. (A, B) dCas9-BFP-KRAB was integrated into human ES or iPS cells either randomly via a PiggyBac construct or specifically into the CLYBL locus by homologous recombination using TALENs. The percentage of BFP-positive cells was monitored over time following neuronal differentiation. (C) EGFP was integrated into the CLYBL locus by gene targeting using CRISPR, and the median EGFP intensity was measured over time following differentiation of ES cells into neurons. (D) Outline of the TK4 PiggyBac vector. (E) EGFP and BFP fluorescence in TK4 vector integrated iPS cells was assessed during neuronal differentiation by flow cytometry at indicated time points. (F) Four different promoters were tested in the TK4 vector by measuring EGFP intensity during neuronal differentiation. Biological replicates did not match most time points and are thus shown separately in Figure S1D. (G) The contribution of the pA signal and the WPRE element to silencing was assessed by EGFP fluorescence intensity during neuronal differentiation by flow cytometry as indicated. The experiments for (E), (G), and (S1D) were done in parallel. For convenience we have re-plotted the EGFP line from (E) as “polyA + WPRE” in (G) and “CAG” in (S1D)”. Biological replicates performed with different time points are shown in Figure S1G.

First, we integrated this TK4 vector by co-transfection with transposase into KOLF2.1 iPS cells that already contained a NGN2-inducible cassette^23^. Upon puromycin selection, cells were differentiated to neurons by addition of doxycyline. After another five days, we added primary mouse glia to support neuronal maturation. We then measured the EGFP fluorescence in neurons over time by flow cytometry. Since mTagBFP2 happened to be fused to our puromycin selection cassette, we also recorded mTagBFP2 fluorescence together with EGFP (**Figures S1A and S1B**). As expected, mTagBFP2 expression was rapidly reduced and barely detectable after 1 week of differentiation compared to undifferentiated iPS cells (**Figure 1E**). Surprisingly, however, the EGFP fluorescence intensity did not decline over time, instead, it rather slightly increased over time (**Figure 1E**). Longitudinal, quantitative fluorescence imaging also confirmed the lack of EGFP silencing in the TK4 PiggyBac vector (**Figure S1C**).

### CAG and Ubc, but not EF1α promoters resist silencing in the TK4 vector

It is long known that the choice of promoters greatly influence the degree of transgene silencing. We therefore tested three widely used promoters that are considered strong promoters in various contexts. CAG, Ubc, EF1α, and EF1α with additional SP1 binding sites (herein referred to as long EF1α) were cloned into the TK4 vector in front of EGFP. We integrated these constructs into iPS cells and differentiated them into neurons with Ngn2. Flow cytometry analysis replicated that the CAG promoter resists silencing of EGFP (**Figures 1F and S1D**). Longitudinal confocal microscopy showed similar results (**Figure S1E**). In addition, the Ubc promoter was comparable to the CAG promoter. In contrast, both EF1α or long EF1α promoters were rapidly silenced. Notably, EGFP expression in undifferentiated stem cells was several fold higher with the EF1α promoters than with CAG or Ubc, but this could not compensate the high degree of silencing after neuronal differentiation (**Figures S1I and J**).

### The Woodchuck Posttranscriptional Regulatory Element is a critical element to prevent silencing

To further assess the contribution of the vector elements to the prevention of silencing of the TK4 vector, we eliminated either the Woodchuck hepatitis virus postranscriptional regulatory element (WPRE)^24^, the polyA signal sequence, or both elements from our PiggyBac vector. The construct without WPRE showed a strong decrease in GFP expression, whereas removing the polyA signal did not significantly affect the expression level during neuronal differentiation (**Figures 1G, S1F and S1G**). In alignment with our previous results, mTagBFP2 expression driven by the EF1α promoter decreased rapidly in all conditions. (**Figure S1H**).

### No major contribution of insulator sequences to prevent silencing resistance

Since our TK4 vector contains several transcriptional insulators, we hypothesized that they may contribute to the prevention of transcriptional silencing. First, we removed all four insulators from our vector, creating a “Null” insulator version of our vector, and compared EGFP expression between the original TK4 vector and the Null vector (**Figure 2A**). Our flow cytometry and longitudinal imaging data showed that the insulators in TK4 vector were not required to prevent silencing unlike the WPRE (**Figures 2B, S2A and S2C**). Unexpectedly, the TK4 vector without any insulator elements performed very similarly to the original TK4 vector.

**Figure 2.**
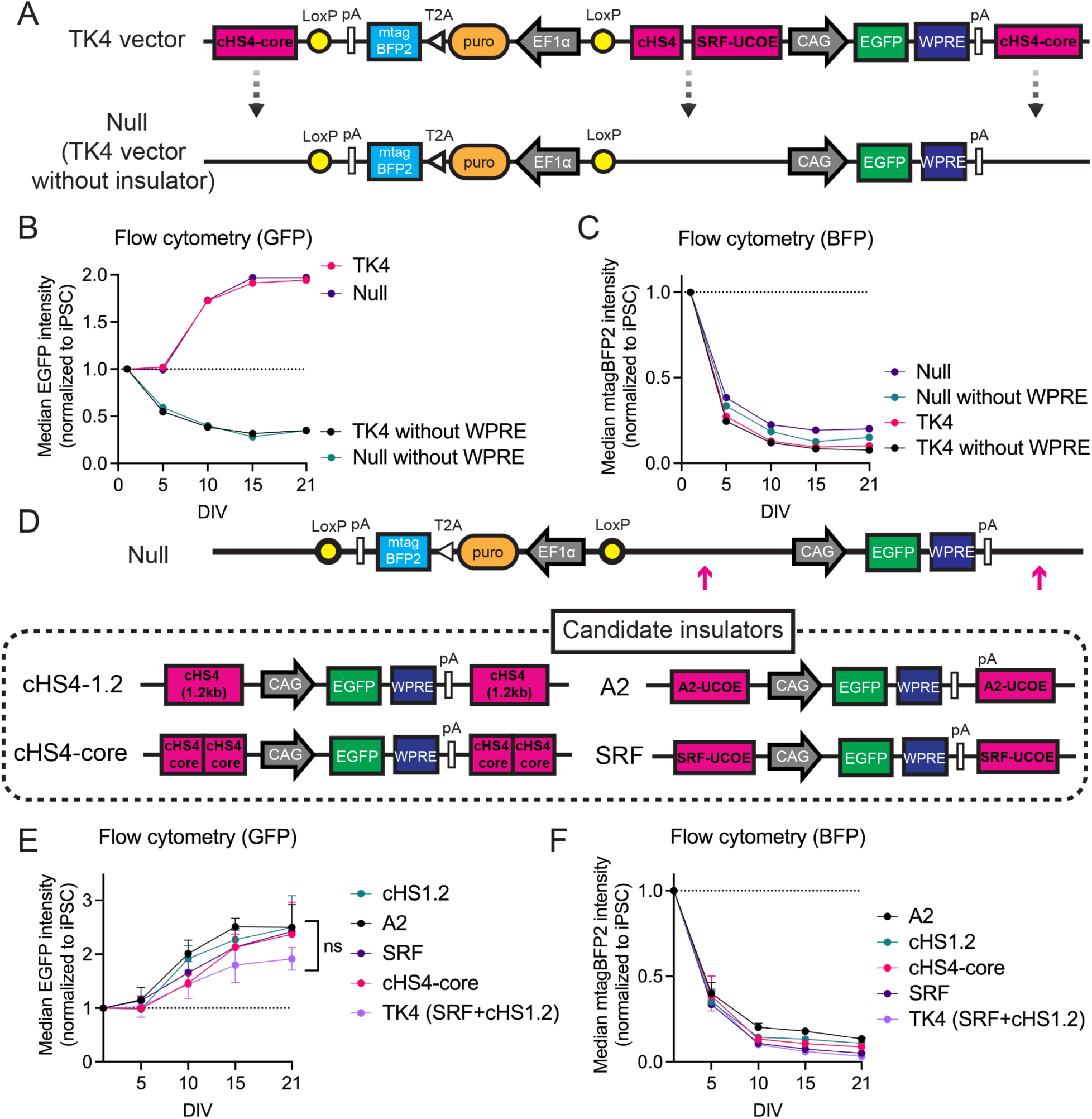
No major impact of insulator sequences on maintaining gene expression from the TK4 vector during differentiation. (A) The insulator sequences were eliminated from the TK4 vector to generate the TK4 Null vector. (B) Median EGFP intensity measured by flow cytometry for each construct, with or without insulators and with or without WPRE, following neuronal differentiation. Biological replicates are shown in Figure S2A. (C) Median BFP intensity measurements corresponding to EGFP measurements shown in (B). Biological replicates are shown in Figure S2B. (D) Candidate insulators were inserted into the Null line at the position indicated. (E) Median EGFP intensity showed no significant difference between constructs with different insulator elements during neuronal differentiation. Statistical analysis was performed using two-way ANOVA (n = 3). (F) As expected, BFP was silenced in all vectors tested.

Nevertheless, we wondered whether different variations of insulator additions may further improve the performance of the TK4 vector. We selected four insulator elements: the full-length 1.2kb long chicken hypersensitive site 4 element (referred to as cHS1.2)^15^, the core of cHS4 (cHS4-core)^25^, A2-ubiquitous chromatin opening elements (A2-UCOE)^5^, and SRF-UCOE^22^. cHS4 acts as an insulator with specific protein binding sites to block heterochromatin spread, while UCOEs use housekeeping gene promoters to create ubiquitously active chromatin regions. We compared these insulators in different configurations, including flanking, upstream, and in the opposite orientation, using confocal microscopy (**Figure S2D**). However, no significant differences were observed between these configurations (**Figures S2E and S2F**). Next, we chose four constructs that trended to perform better than the original TK4 vector and tested them by flow cytometry. These constructs contained the four insulators in a flanking position (**Figure 2D**). Again, there was a trend that these constructs lead to higher expression levels than the original TK4 vector upon differentiation but the differences were not statistically significant (**Figure 2E**). As expected, mTagBFP2 intensity in these lines that contained these various insulators showed consistent silencing during differentiation as measured by flow cytometry (**Figures 2C, 2F, and S2B**). These data suggest that insulator elements may not be as critical as promoters or the WPRE for stable transgene expression during neuronal differentiation.

### The TK4 vector prevents silencing of non-EGFP transgenes

For this experiment, we wanted to test another reporter gene in order to ensure that the effects observed so far are not unique to EGFP. We decided to use the red fluorescent protein mScarlet followed a destabilized mNeonGreen protein joined with a T2A sequence that is predicted to produce two separate proteins. The inclusion of the destabilized mNeonGreen was an attempt to obtain a more sensitive expression readout than with stable fluorescent proteins. However, upon integration into iPS cells we could not detect mNeonGreen expression in undifferentiated or differentiated cells and therefore focused only on measuring mScarlet for the quantification of transgene silencing.

For best comparison with the EGFP results, we first cloned the mScarlet-mNeonGreen expression cassette into the TK4 PiggyBac vector (**Figure S3A**). Flow cytometry confirmed that mScarlet fluorescence rather increased than decreased during neuronal differentiation, just like we had observed for EGFP as readout before (**Figure 3A**).

**Figure 3.**
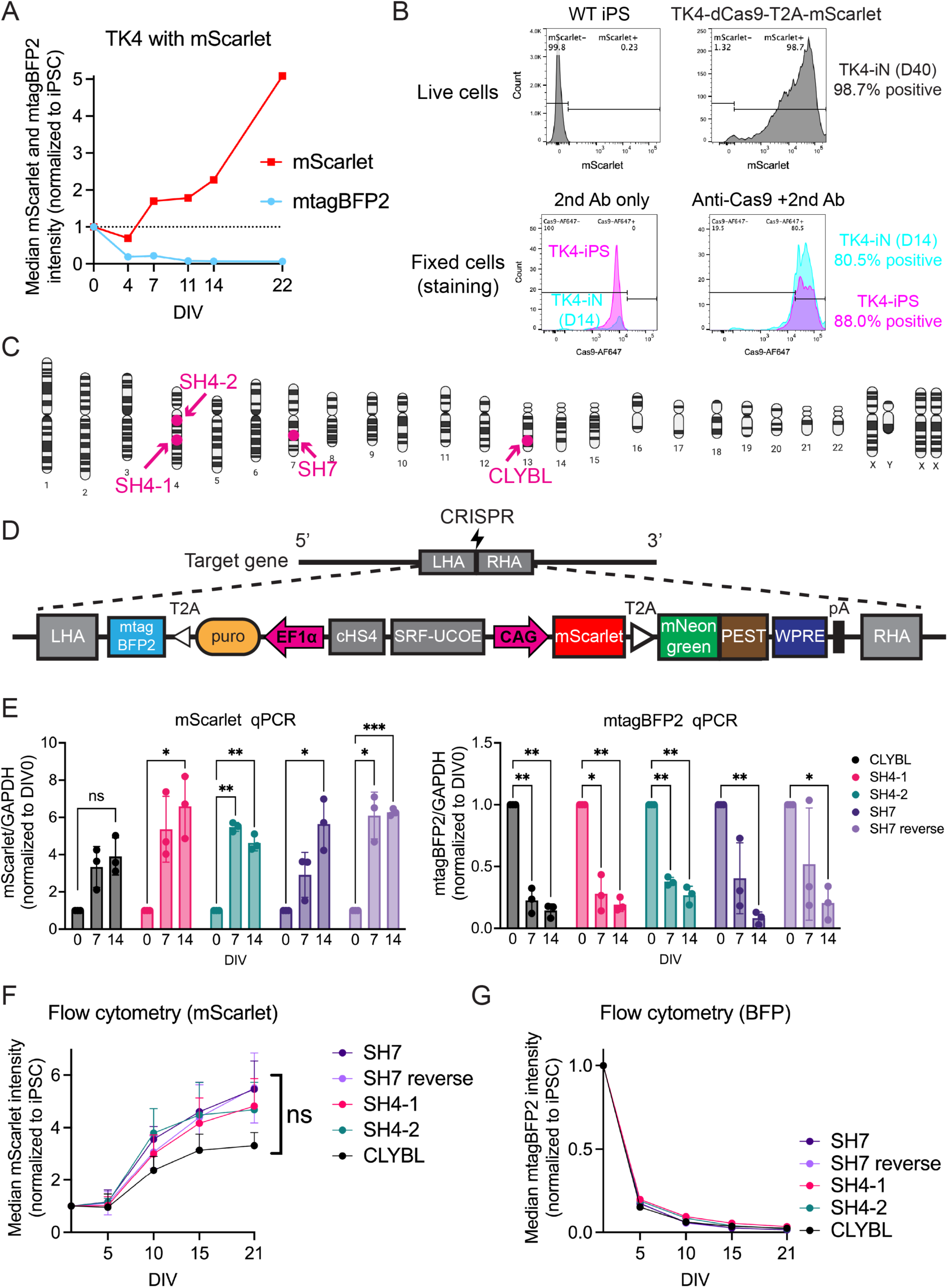
Three new safe harbor loci for stable transgene expression of the TK4 expression cassette. (A) As seen for EGFP before, mScarlet fluorescence is not silenced in the TK4 during neuronal differentiation of iPS cells. (B) Top histogram: Robust mScarlet expression in neurons 40 days after differentiation. Bottom histogram: Comparable and strong expression of Cas9 as detected by immuno flow analysis of undifferentiated iPS cells and day 14 neurons. (C) The *CLYBL* and three novel safe harbor loci, distant from annotated genes, were selected for targeted integration of the TK4 expression cassette^12^. (D) The expression construct of the TK4 vector was placed between targeting homology arms for CRISPR-mediated homologous recombination. Note: the destabalized mNeonGreen:PEST fluorescence was not detected by flow cytometry or microscopy and therefore not further evaluated. (E) qRT-PCR analysis of mScarlet and mTagBFP2 at day 0 (iPS cells), day 7, and day 14 of neuronal differentiation. *P < 0.05, **P < 0.01, ***P < 0.001; Two-way ANOVA with Dunnett’s multiple comparisons test. n = 3. (F) Flow cytometry was used to measure the median mScarlet intensity during neuronal differentiation, normalized to the intensity in undifferentiated iPS cells for each targeted line. Statistical analysis was conducted using two-way ANOVA. n = 3. (G) BFP intensity in the indicated lines, equivalent to the analysis in (F). n = 3.

Next, we wondered how a transgene coding for a non-fluorescent protein would perform in the TK4 vector. To this end, we cloned the commonly used CRISPR interference reagent dead Cas9-Zim3 tagged with mScarlet into the TK4 PiggyBac vector (**Figure S3B**). At day 14 and 40 of neuronal differentiation, we confirmed that almost all neurons exhibted mScarlet fluorescence (**Figure 3B, top and Figure S3B, right**). Furthermore, we also stained neurons at day 14 with anti-dCas9 and anti-HA antibodies to detect the HA-tagged dCas9 construct. Flow cytometry demonstrated that dCas9 and HA intensity were similar between differentiated neurons and undifferentiated iPS cells (**Figure 3B, bottom and Figure S3B, left**).

These results demonstrate that the lack of silencing in the TK4 vector is not due to the EGFP reporter.

### Three novel genomic loci for stable transgene expression

Next, we wanted to compare the performance of the expression cassette in the TK4 vector in various defined genomic loci in hopes to identify a locus that may outperform popular integration sites such as the *CLYBL* locus, which show silencing of transgenes during neuronal differentiation (**Figures 1B and 1C**). Moreover, most current “safe” harbor sites do in fact destroy existing genes. A recent study developed a barcoded synthetic core promoter reporter integrated randomly by PiggyBac to measure the level of transcriptional support across thousands of sites in the human genome during the first five days of neuronal differentiation of iPS cells^12^. Among these characterized integration sites, we chose three that would represent good candidates of safe harbor locations. From the high confidence integration sites reported, we chose three that were (i) in the “on-on group” (group 2 in that study) that were active in both iPSCs and in differentiated neurons, (ii) in intergenic regions and distant from annotated genes, (iii) showed the highest expression levels (maximal cDNA/gDNA ratio). Two of these sites were on chromosomes 4 and one on chromosome 7 (**Figure 3C**).

We identified CRISPR cut sites in the vicinity of these loci and generated targeting constructs by amplifying and cloning homology arms for these 3 loci into the TK4 vector (**Figures 3D, S3C, and S3D**). Transient nucleofection of guideRNA/Cas9 ribonuclear protein (RNP) complexes facilitated the homologous recombination of expression constructs. For the chromosome 7 integration site we generated the knock-ins in both orientations. After puromycin drug selection iPS cell clones were picked and genotyped for proper transgene integration.

Next, we assessed mScarlet expression in five iPS cells lines with mScarlet targeted to the two sites on chromosome 4, the site on chromosome 7 in both orientations, and the *CLYBL* locus as comparison. qRT-PCR analysis showed that mScarlet mRNA even increased following neuronal differentiation across all loci tested. In contrast and as expected, mTagBFP2 mRNA expression decreased progressively over time (**Figure 3E**). Consistent with the mRNA expression results, mScarlet fluorescence measured by flow cytometry showed an increase over time following neuronal differentiation (**Figure 3F**), while mTagBFP2 expression was rapidly silenced (**Figure 3G**). Compared to the *CLYBL* locus, our candidate loci showed a stronger and significant increase over time at the RNA level and a non-significant increase on the protein level. Especially both orientations of the transgene integration at chromosome 7 resulted in non-significant but consistently higher mScarlet fluorescence among three biological replicates than the *CLYBL* integration site.

These data illustrate that our three novel safe harbor loci serve as excellent expression loci at least on par with the *CLYBL* locus. In all loci the expression elements of the TK4 vector prevented transgene silencing. With regards to overall expression levels, the three novel loci slightly outperformed the *CLYBL* locus at least at the RNA level. In addition, all three of our new loci are outside of any annotated transcriptional units, offering safer options for defined genomic integrations than currently used loci, especially in a therapeutic context.

### TK4 vector resists silencing in various neuronal subtypes cells and microglia derived from iPS cells as assessed by seven different laboratories

Finally, we wanted to explore whether our results could be applied to additional cell types and differentiation protocols and are consistent between different laboratories. To address this, we shipped our TK4 PiggyBac vector to six additional labs in the US and Europe (**Figure 4A**). Researchers independently created TK4-integrated human iPS cells in 1 or 2 different cell lines and differentiated them into Ngn2-iN cells (**Figure 4B**), transcription factor-programmed microglia^26^ (**Figure 4C**), dopaminergic neurons^27^ (**Figure 4D**), cortical neurons^28, 29^ (**Figure 4E**), microglia using an embryoid body-based protocol^30^ (**Figure 4F**), microglia using a monolayer-based protocol^31^ (**Figure 4F**), striatal neurons^32^ (**Figure 4G**), and hypothalamic neurons^33^ (**Figure 4H**). Each group independently measured transgene expression over time during differentiation by flow cytometry as described above. Remarkably, without exception, the TK4 vector consistently showed stable or increased EGFP expression over time, while mTagBFP2 was silenced, in aggregate in six different human iPS cell lines and nine different protocols. We conclude that the stable gene expression of the TK4 vector during neuronal and microglial differentiation of human iPS cells is robust and highly reproducible.

**Figure 4.**
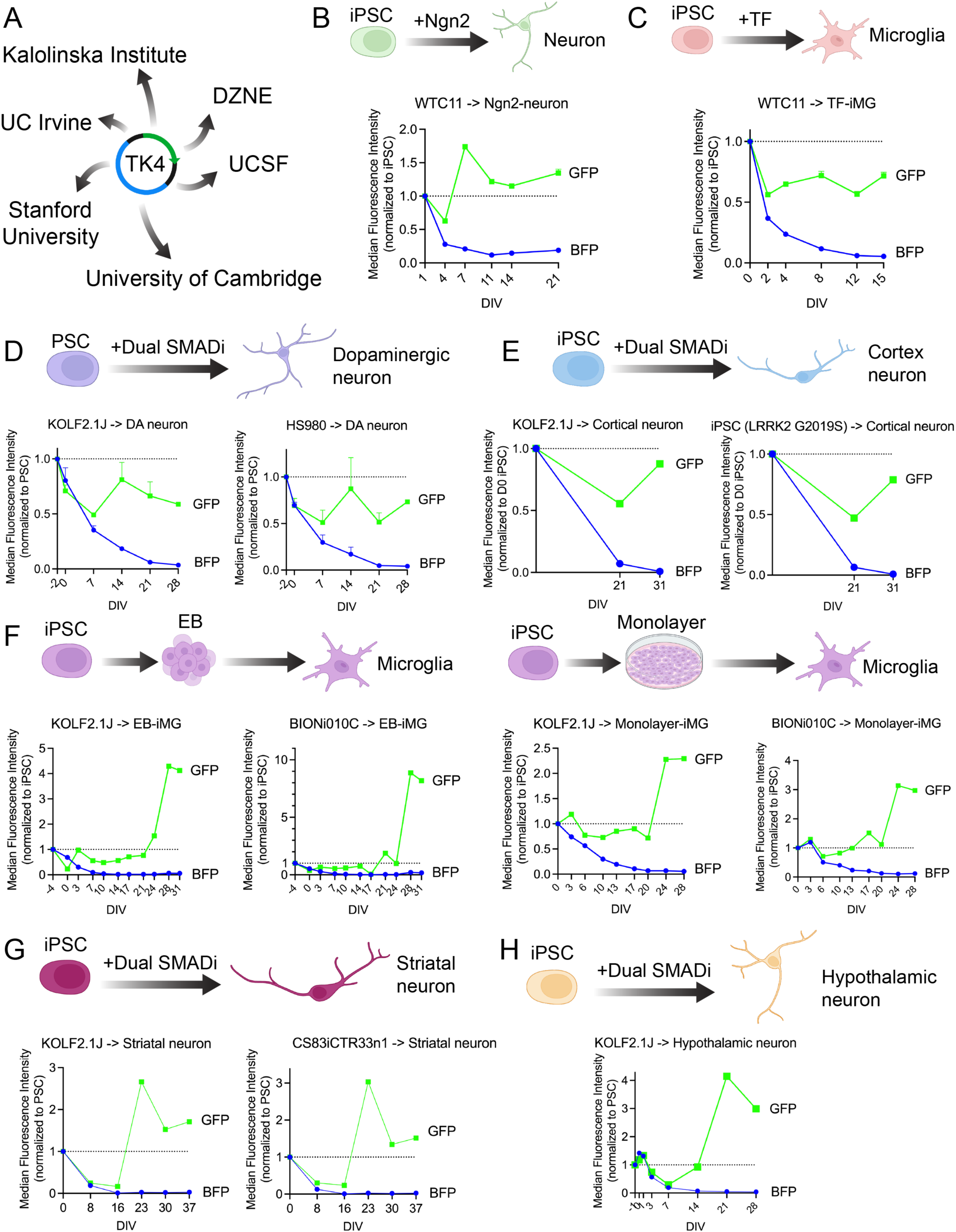
The TK4 vector resists silencing among various neuronal and microglial differentiation protocols. (A) The TK4 vector was distributed to 6 different laboratories. Each lab transfected the vector into various iPS cell lines and performed differentiation into neurons or microglia using different protocols. EGFP and BFP fluorescence was measured by flow cytometry and normalized to the expression level of undifferentiated iPS cells. (B) Ngn2-neuronal mono-culture from the WTC11 iPS cell line. n = 1 with 6 technical replicates. (C) Transcription factor-mediated differentiation to microglia from the WTC11 line. n = 1 with 3 technical replicates. (D) Dopaminergic neurons from KOLF2.1J and HS980 stem cell lines. n = 2. (E) Cortical neurons from 2 different human iPS cell lines. n = 1. (F) Microglia derived through an embryoid body (EB)-containing and a monolayer differentiation protocol from KOLF2.1J and BIONi010C iPS cell lines. n = 1. (G) Striatal neurons generated from KOLF2.1J and CS83iCTR33n1 iPS cell lines. n = 1. (H) Hypothalamic neurons from KOLF2.1J line. n = 1 with 3 technical replicates.

## DISCUSSION

We have established a PiggyBac system that effectively prevents transgene silencing during the differentiation of neurons or microglia from human stem cells. This construct has also proven useful for targeting specific loci, with reproducible results across various differentiation protocols conducted by different laboratories.

The silencing of experimental transgenes during differentiation of human pluripotent stem cells is a not much publicized but widely-known problem in stem cell biology labs and unfortunately often re-discovered by scientists new to the field. Even simple genetic approaches, like expression of disease genes or genetic rescue of gene knock-out experiments often become unfeasible, let alone the use of sensors such as voltage or Ca^2+^ indicators or Cas9-related enzymes for gene manipulation. Choosing strong constitutive promoters, targeting defined expression loci, or use of insulator elements have resulted in highly stable transgene expression in pluripotent stem cells as long as they remain undifferentiated^34, 35, 36^. Remarkably, though, transgenic silencing will invariably occur during differentiation into somatic cell types^37, 4, 38^. At this point, the mechanisms of transgene silencing during pluripotent cell differentiation are not understood. Pluripotent cells are in a unique epigenetic state. E.g. they contain bivalent histone domains and in contrast to somatic cells, they can tolerate complete loss of DNA methylation, loss of histone methylases like G9a, and loss of the RNAi machinery^39, 40, 41, 42, 43^. Of note, γ-retroviruses are acutely silenced in pluripotent and early embryonic cells and the silencing is mediated by DNA methylation^44, 45^. Perhaps when cells leave this unique epigenetic state as they exit pluripotency, decisive epigenetic changes need to be set in motion that overwhelm experimental expression cassettes, no matter where they are integrated.

Silencing has been mostly studied in highly proliferative cell lines. Like in pluripotent cells, epigenetic mechanisms including DNA and histone methylation have been implicated along with cell cycle-related mechanisms like R-loop formation, engagement of viral and transposon responses, and spreading of nearby heterochromatin^37^. Since cell proliferation rather decreases during neuronal differentiation it is unlikely that cell cycle-related mechanisms are relevant. We observed no silencing of our randomly integrating PiggyBac vector and in four different genomic loci. Moreover, various insulator sequences tested had little effect on transgene silencing. Hence, positional variegation effects such as heterochromatin spreading may also be of less importance in our system. Instead, we found that promoter sequences and a posttranscriptional regulatory element were primarily responsible for the lack of silencing of the TK4 vector. The sequences of promoters are known to influence gene silencing. In particular viral elements such as CMV, SFFV, and γ-retroviral LTR promoters are subject to substantial silencing in mammalian cells^46, 47^. However, even mammalian promoters that express well and stably in undifferentiated cells such as EF1α or Ubc are silenced in a conventional expression configuration upon differentiation (while obviously their physiological genes are not). Remarkably, however, a subset of these stable promoters will resist silencing if the expression cassette is combined with a Woodchuck hepatitis virus-derived WPRE. The molecular mechanisms of how the WPRE increases cytoplasmatic mRNA are poorly understood. Based on sequence similarity to the Hepatitis B Virus PRE it is assumed that it enhances nuclear export of mRNA^48^. Other potential mechanisms proposed include the support of transcriptional termination and 3’ RNA processing^24, 49^. Since all of these mechanisms are promoter independent, our observation that the WPRE only blocks silencing when combined with certain promoters suggests the existence of yet additional mechanisms for WPRE’s function. Perhaps it interacts with the 5’ end of the transcribed RNA molecule such as transcriptional looping which can result in transcriptional stabilization and depends on promoter sequences^50^.

For both experimental and therapeutic applications, there is much debate about the characteristics of an optimal “safe harbor” site in the genome that can be used to express transgenes^51^. Due to unpredictable effects on transgenes targeted to gene deserts, the field has mostly used integrations into human genes considered less important (such as *CLYBL*, *AAVS1*, *ROSA26*, and *CCR5* loci). These sites may be safe to ensure expression but they are certainly not perfectly safe from a perspective for minimal interference with the host genome as they all disrupt endogenous human genes. In this study, we took advantage of a recently developed resource that mapped genome wide integration sites with level of gene expression^12^. We purposely chose three novel candidate integrations that were distant from any annotated human genes and therefore have a greater potential to be actual safe harbor sites. All three of these novel expression loci performed as well as the *CLYBL* locus in terms of expression levels, if not even better. If gene disruption of the conventional safe harbor sites is of concern, these new sites could be an attractive alternative.

In summary, we here have developed a silencing-resistant expression construct that we hope will have a major impact on the field as it will enable experiments that require robust transgene expression in somatic cells differentiated from pluripotent stem cells.

## Limitations

The main limitation of this study is that we currently do not know whether the TK4 vector will prevent silencing in all somatic cells that can be generated from human pluripotent stem cells. We are quite confident about neuronal and myeloid cell types because those were the main focus of this study. We hope our vector will be tested by more scientists in the stem cell field to address this important question.

Another critical limitation of this study is the number of reporter genes tested. Most of our results are based on EGFP expression. In addition, we have confirmed that the red fluorescent protein mScarlet and dCas9 behaves like EGFP. However, it has been seen before that the transgene used affects the degree of silencing. It is therefore well possible that the TK4 will not be able to resist silencing of all possible transgenes.

Finally, the strong promoter dependence we observed will require to test any additional promoters that may be desired such as cell-type specific or inducible promoters.

## ACKNOWLEDGMENTS

This work was conceptualized by the Chan-Zuckerberg-Initiative Neurodegeneration Challenge Network’s Stem Cell Working Group led by Margaret Sutherland and supported by a grant from the Chan-Zuckerberg-Initiative. We would like to thank the FACS core at the Institute for Stem Cell Biology and Regenerative Medicine of Stanford University and Wu Tsai Neuroscience Institute Neuroscience Microscopy Service of Stanford University for their microscopy assistance, expertise, and services, Gang Ning for help with genotyping of clonal knockin cells and FACS, Lacramioara Bintu and Michael Bassik for critical comments on the manuscript and Alaka Mullick together with all members of the working group for helpful discussions and comments on study design and execution of this project. Hsiao-Jou Cortina Chen, Eugene Seah and Amber Keith provided technical support. T.U. was supported by the Wu Tsai Neurosciences Institute Knight Initiative for Brain Resilience Scholar Award. A.M. was supported by Postdoctoral fellowships from the Alzheimer’s Association (AARF-22-973222) and the Larry L. Hillblom Foundation (2022-A-016-FEL). A.J.S was supported by NIH/NIA Pathway to Independence Award (K99AG080116). L.E. was supported by the Studienstiftung des Deutschen Volkes (German Academic Scholarship Foundation, PhD Scholarship) and the International Max-Planck Research School for the Mechanisms of The Mental Function and Dysfunction (IMPRS-MMFD, PhD Scholarship). M.K. was independently supported by the Chan Zuckerberg Initiative (2022-316571). L.M.T was supported by NIH grants (R35NS116872).

## AUTHOR CONTRIBUTIONS

M.W. and M.E.W. conceived the project. M.W., M.E.W., and T.U. designed the experiments. T.U., A.B.N., A.D.S., D.S., A.S., L.C., J.C., L.E., A.M., D.R., L.S., A.J.S., K.S., K.W., and N.W. performed the experiments. T.U., A.B.N., A.D.S., A.S., L.C., J.C., L.E., A.M., D.R., L.S., A.J.S., K.S., K.W., N.W., and Q.W. developed experimental protocols, tools, and reagents or analyzed data. W.C.S. generated the safe-harbor lines. M.W., T.U., J.C., L.E., A.M., L.S., A.J.S., K.S., N.W., E.A., A.R.B., M.K., D.K.-V., F.T.M., B.S., L.M.T., W.C.S., and M.E.W. wrote the manuscript.

---

**Figure S1.**
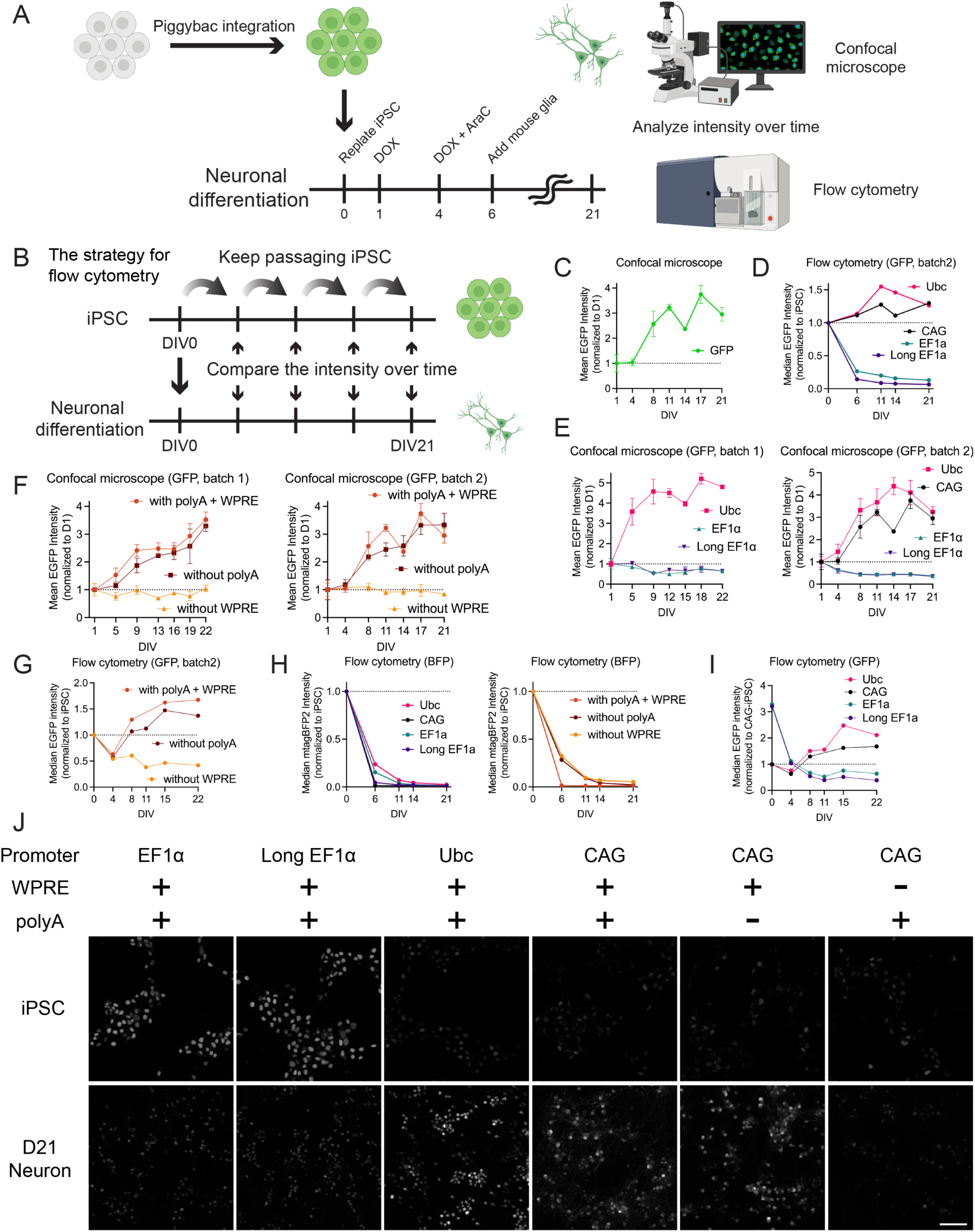
Comparison of Promoters and Regulatory Elements in the TK4 Vector. (A) Schematic diagram depicting the design of this project. (B) Schematic diagram illustrating the process for measuring fluorescence by flow cytometry during neuronal differentiation. (C) Mean EGFP intensity measured by confocal microscopy in TK4 line. Data consists of technical triplicates. (D) Additional biological replicate of flow cytometry corresponding to Figure 1F . (E) Mean EGFP intensity measured by confocal microscopy in each line, comparing different promoters. Two biological replicates are shown with technical triplicates in each plot. (F) Mean EGFP intensity measured by confocal microscopy in each line, comparing different regulatory elements. Two biological replicates are shown with technical triplicates in each plot. (G) Additional biological replicate of flow cytometry corresponding to Figure 1G. (H) Median BFP intensity following neuronal differentiation among different promoters and regulatory elements. The line with “CAG” was common with the “polyA + WPRE” line. (I) Median EGFP intensity measured by flow cytometry and normalized to iPS line with CAG promoter. (J) Representative confocal microscope images of iPS cells and day 21 neurons. Scale bar, 100 μm.

**Figure S2.**
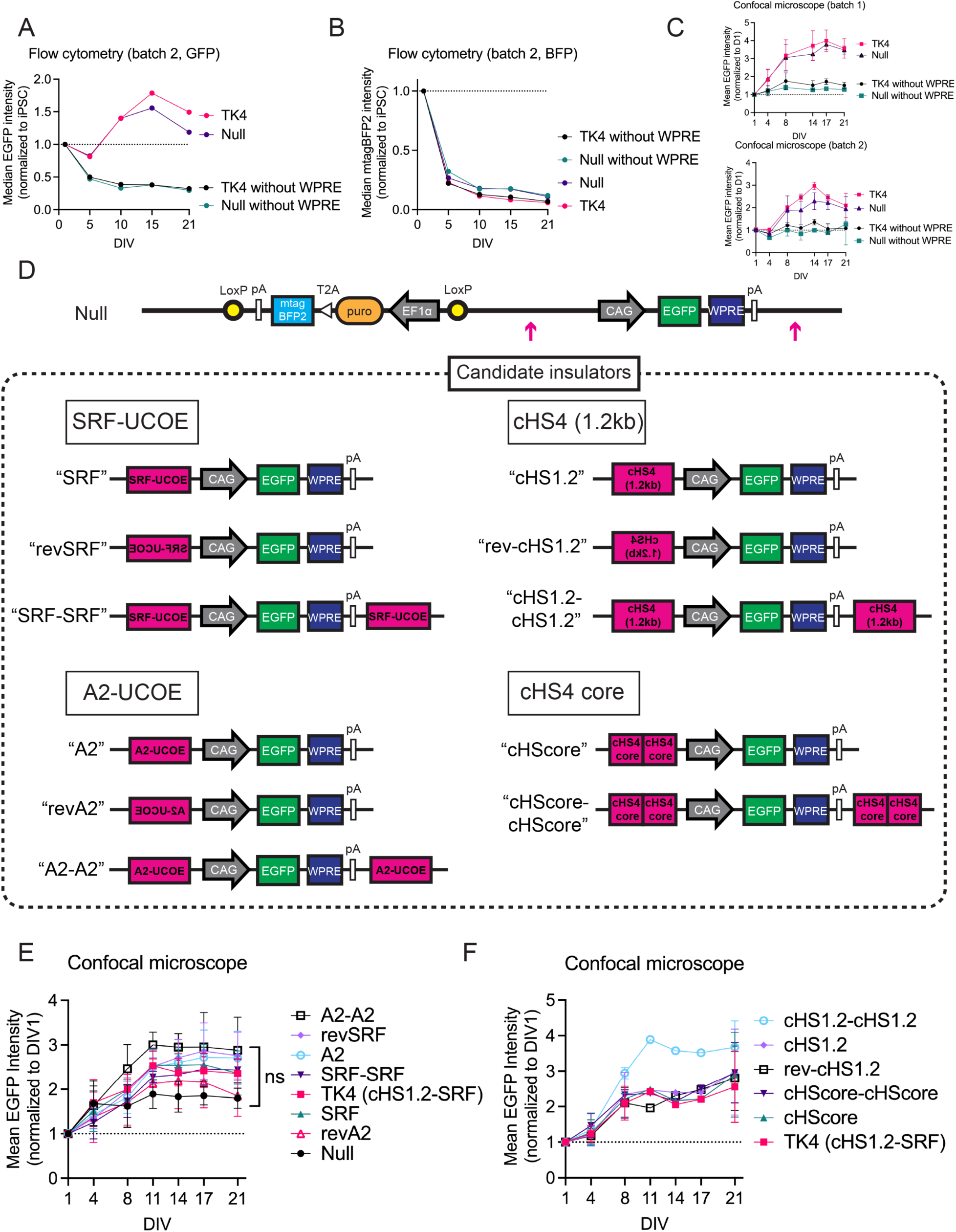
Comparison of different insulators in the TK4 vector. (A) Additional biological replicate corresponding to Figure 2B. (B) Additional biological replicate corresponding to Figure 2C. (C) Mean EGFP intensity measured by confocal microscopy in each line, comparing with or without insulator elements and with or without WPRE. Two biological replicates are shown with technical triplicates in each plot. (D) Candidate insulators were inserted into the Null line at the position indicated by the arrow. (E) Mean GFP intensity was assessed by confocal microscopy following neuronal differentiation across different insulator lines. Statistical analysis was performed using two-way ANOVA. n = 5. (F) Mean GFP intensity was assessed by confocal microscopy following neuronal differentiation across different insulator lines. n = 2.

**Figure S3.**
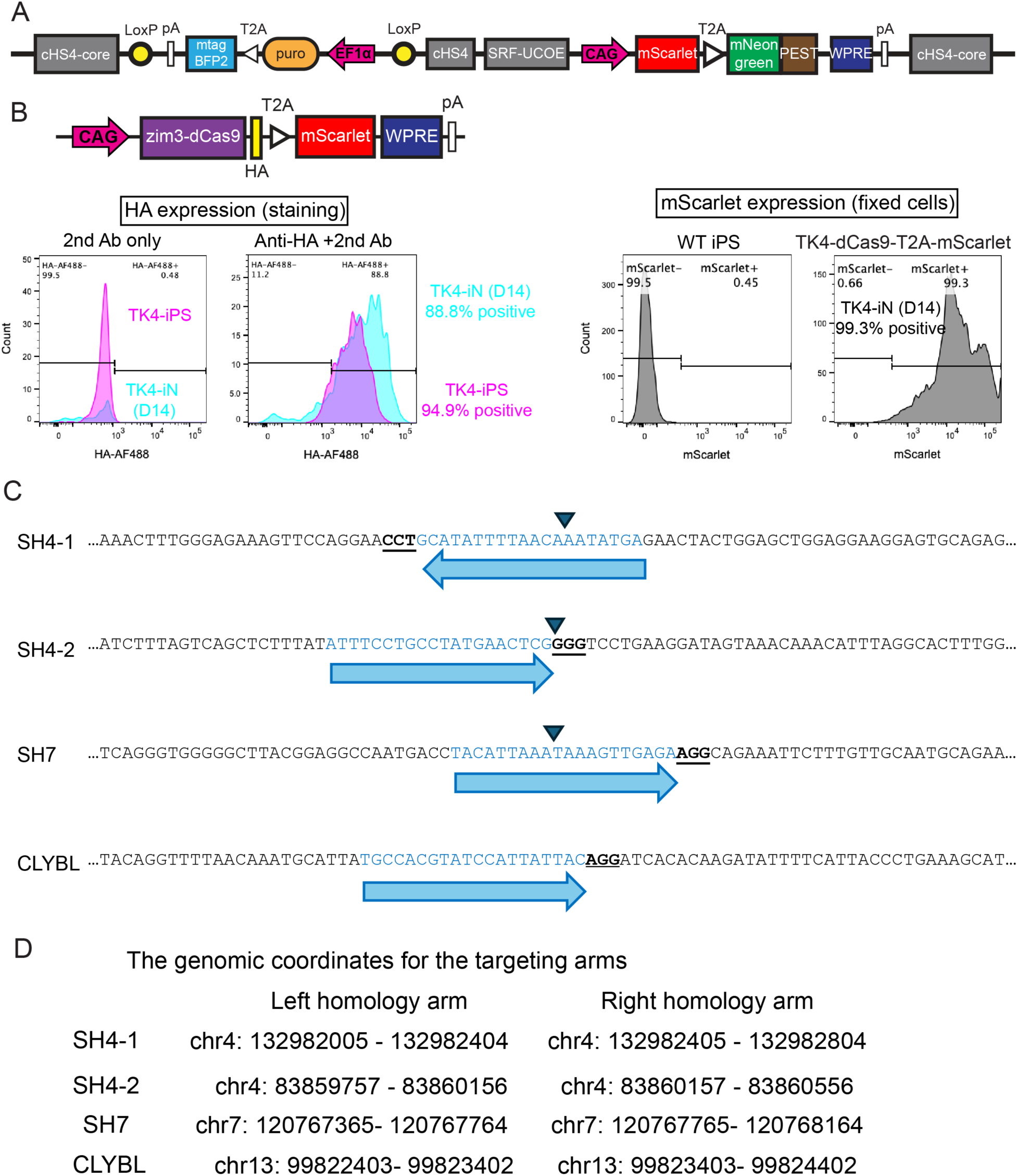
Knock-in of TK4 transgene into new candidate safe-harbor loci. (A) Schematic of the mScarlet-T2A-mNeonGreen:PEST expressing TK4 vector. Note: the destabilized mNeonGreen:PEST protein was not detectable by flow cytometry or microscopy and hence not further evaluated. (B) Schematic of the Zim3-dCas9-HA-T2A-mScarlet TK4 vector. Left histogram: Stable expression of HA, which is fused with dCas9, as detected by immune flow analysis of undifferentiated iPS cells and day 14 neurons. Right histogram: Robust expression of mScarlet in day 14 neurons. (C) DNA sequences for each safe harbor locus and corresponding sgRNAs. Triangles indicate the PiggyBac insertion sites identified by Wu et al^12^. (D) The genomic coordinates (hg38) for the targeting arms in each locus.

**Figure S4.**
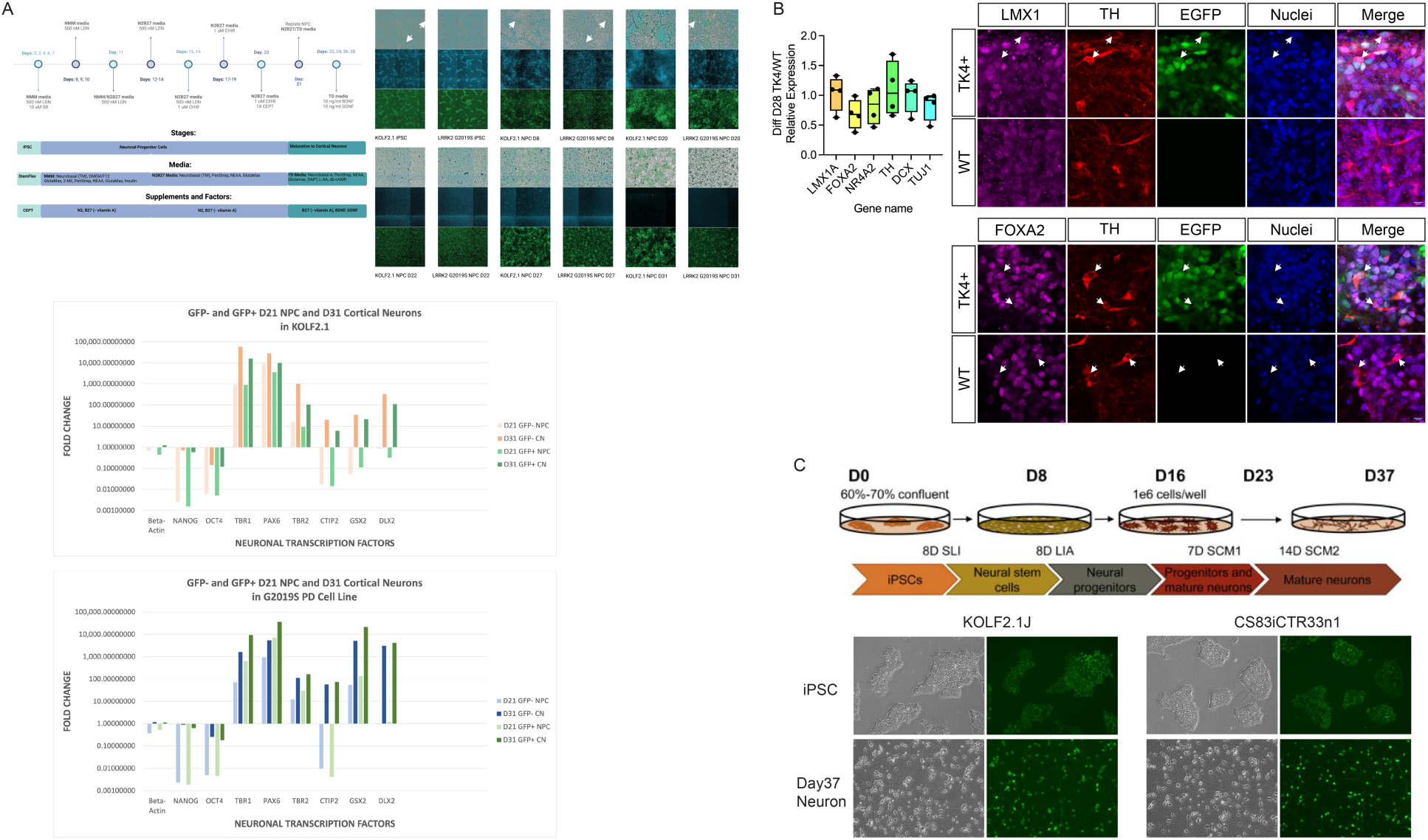
Characterization of various neuronal differentiation protocols. (A) (top) Schematic of the differentiation from iPS cells to cortical neurons and the signal of BFP/EGFP during differentiation. (bottom) mRNA expression in D21 and D31 differentiaion with or without GFP.Verification of dopaminergic neuron differentiation capacity in TK4-EGFP reporter HS980 and KOLF2.1J pluripotent stem cell lines. Human iPSC line KOLF2.1J and human ESC line HS980 stably expressing the TK4 reporter were differentiated to midbrain dopaminergic neurons for 28 days. (Left) Expression of marker genes in GFP-expressing cells relative to untransfected WT parental controls at d28 differentiation (Kolf2.1J n = 2; HS980 n = 2). (Right) Immunocytochemistry of LMX1 and FOXA2 floorplate markers, TH dopaminergic marker and EGFP expression after 28 days differentiation of KOLF2.1J cells towards a mDA lineage. Arrows indicate examples of quadruple positive cells. Nuclei (Dapi, blue). Scale bars are 10 μm. Stable expression of TK4-EGFP reporter does not impair the ability of hPSCs to be differentiated towards a dopaminergic lineage. (B) (top) The scheme of differentiation from iPS cells to striatal neurons. (bottom) Representative images of phase contrast and GFP in undifferentiated cells and day 37 striatal neuron in 2 different iPS cell lines. The images are taken on the EVOS M5000 with 10X magnification.

## METHOD DETAILS

### iPS Cell Culture and Generation of Polyclonal Lines Using PiggyBac Vector (Marius Wernig lab)

Male human embryonic stem cells (ESCs) line H1 were obtained from WiCell Research Institute. KOLF2.1J iPS cell line, containing TetO-Ngn2 and CAG-rtTA (obtained from Jackson Laboratory) and cultured on Matrigel-coated plates in StemFlex media. For transfection, 1.5 × 10⁶ iPSCs per well were seeded in a 6-well plate with 1.5 mL of E8 media supplemented with Y-27632 and incubated for 2 hours. The transfection mix was prepared by combining 1333 ng of the TK4 vector with 666 ng of transposase in 250 μL OPTI-MEM, along with 5 μL of Lipofectamine STEM reagent. This mixture was incubated at room temperature for 30 minutes. After the 2-hour incubation, wells were washed with DPBS, followed by the addition of 1.5 mL of E8 media with Y-27632. The transfection mix was then added dropwise to the cells. The following day, the medium was replaced with StemFlex media containing Y-27632. Two days post-transfection, Y-27632 was removed, and three days post-transfection, puromycin selection (2 μg/mL) was started. The cells were passaged at least twice in puromycin-containing media.

### Generation of Ngn2-Induced Neurons (Marius Wernig lab)

iPSCs were differentiated with slight modifications to a previously described protocol^52^. KOLF2.1J-Ngn2 cells were plated as dissociated cells in a 24-well plate (10–20 × 10³ cells) or 384-well plate (1–2 × 10³ cells) on day 0. H1 cells without Ngn2 cassette were plated with lentiviruses containing expression constructs of Ngn2-T2A-puromycin and rtTA. On day 1, the culture medium was replaced with N3 media (composed of DMEM/F12 [Thermo Fisher], N2 supplement [Thermo Fisher], and MEM Non-Essential Amino Acids [Thermo Fisher], supplemented with insulin [Sigma]), containing doxycycline (2 μg/mL, Sigma) to induce Ngn2 expression. On days 4 and 5, AraC (4 μM, Sigma) was added to eliminate dividing cells. On day 6, passage 3 mouse glial cells were added on top of the neurons (3 × 10⁴ mouse glial cells per well of a 24-well plate, and 1.5 × 10³ cells per well in a 384-well plate), re-suspended in Neurobasal media (Thermo Fisher) supplemented with B27 (Thermo Fisher), Glutamax (Thermo Fisher), and 5% fetal bovine serum (FBS, GE Healthcare). On day 7, the medium was changed to Neurobasal media with B27, Glutamax, 2% FBS, and AraC (4 μM). Approximately 30% of the media was exchanged every 3-4 days. All animal studies were approved by the Administrative Panel on Laboratory Animal Care (APLAC, Stanford University), and CD1 mice (Charles River) were used in this study.

### Flow cytometry analysis (all participating labs)

iPS cells containing each PiggyBac vector were prepared in 24-well plates as described above. At each time point, the cells were dissociated using 250 μL of Accutase, resuspended in FACS buffer (R&D) supplemented with Y-27632, and filtered through a 70 μm strainer for flow cytometry analysis using a FACSAria instrument. After doublet discrimination, the median fluorescence intensity was measured using FlowJo software.

### Confocal microscopy analysis (Marius Wernig lab)

iPS cells containing each PiggyBac vector were prepared in 384-well plates as described above. At each time point, images were captured using a Nikon Spinning Disk confocal microscope (Nikon) or LSM980 (Zeiss) with a 20x objective lens, utilizing 4-tile imaging. GFP-positive areas were identified, and samples that were too sparse or too dense were excluded based on the area count. EGFP intensity within the positive areas was quantified using ImageJ software.

### iPS cell culture and nucleofections for gene targeting (Bill Skarnes lab)

KOLF2.1J cells^27^ were cultured on Synthemax or vitronectin-coated plates in complete StemFlex media and nucleofected with Cas9 RNP as described previously^53^. TK4 construct with 400 bp homology arms was integrated into 3 safe harbor loci (SH4-1, 4-2 and 7) and 1,000 bp homology arms was integrated into CLYBL locus by CRISPR genome editing. Plasmid DNA donors were ethanol precipitated and resuspended in P3 primary buffer prior to use. Each nucleofection contained two micrograms of plasmid DNA donor, 20 ug HiFi Cas9 recombinant protein (IDT) and 16ug guide RNA (Synthego) in a volume of 0.1 ml Primary buffer 3 plus Supplement (Lonza). Nucleofected cells were plated in Stem Flex media containing ROCK inhibitor (1X Revitacell; Invitrogen). After 48 hours, the media was replaced with Stem Flex containing 0.5 microgram/ml puromycin and changed every two days until colonies formed. The colonies were pooled and cultured in presence of puromycin for two passages prior to single cell cloning. Four hundred single cells were plated on a vitronectin-coated 10cm dish in Stem Flex media containing CloneR2 (Stem Cell Technologies) and the media was changed every two days until colonies appeared. Individual colonies were picked into replicate vitronectin-coated 96-well plates, expanded and archived or lysed for genotyping as described^53^.

### Genotyping (Bill Skarnes lab)

PCR amplification and Sanger sequencing was performed from diluted crude whole cell lysates^53^. Briefly, lysates were diluted 1:10 and 2ul was used in a 20 microliter reaction containing PrimeSTAR GXL polymerase (Takara) and 0.25 micromolar PCR primer pairs spanning the 5’ and 3’ homology arms. Correctly targeted clones were identified by gel electrophoresis and GFP expression was confirmed by FACS.

**Table 1.**
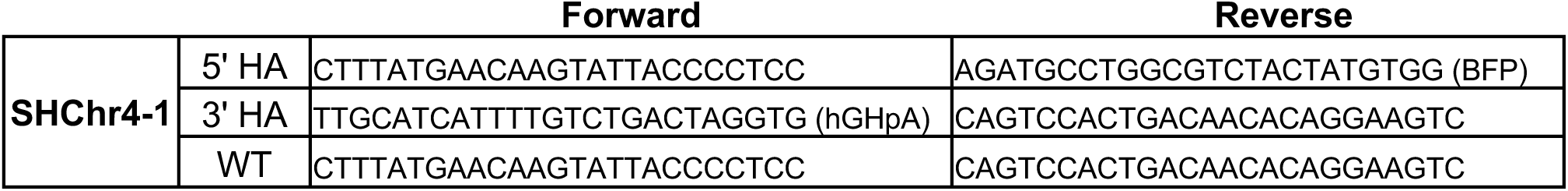
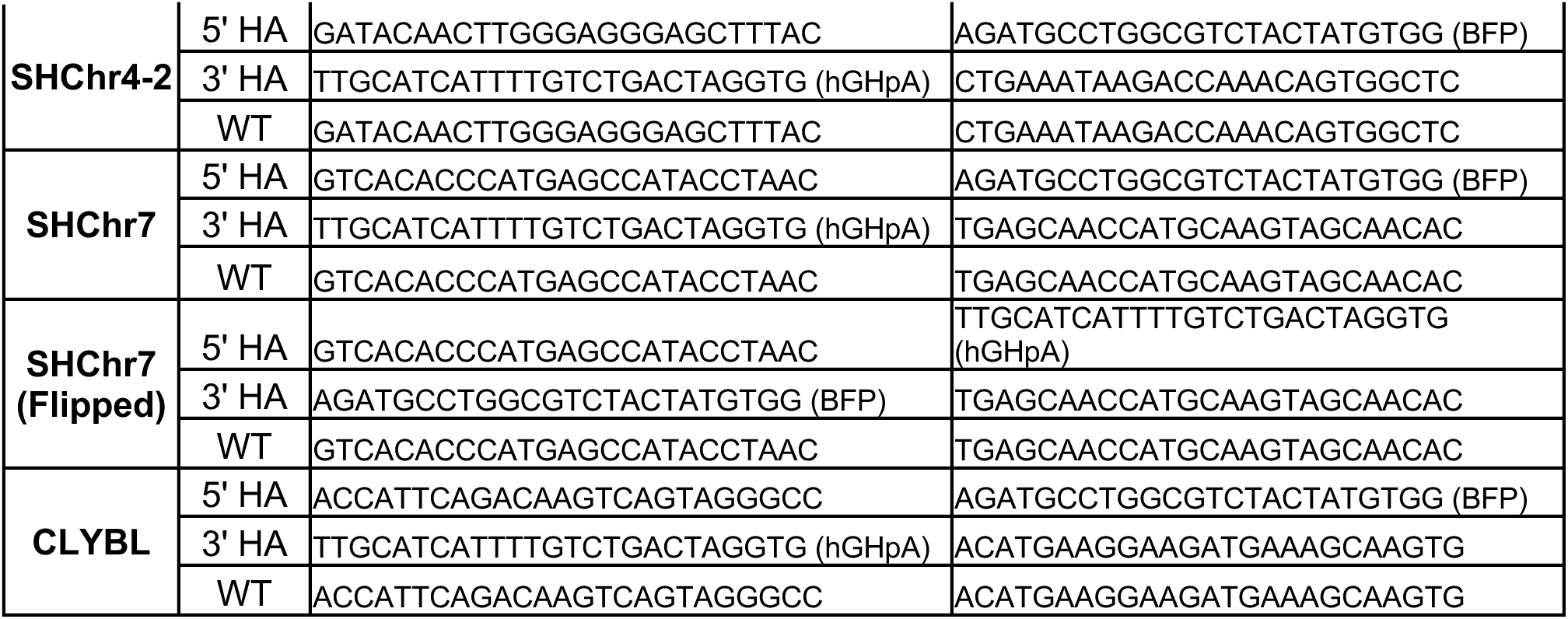
PCR primers for genotyping knock-in clones. Primers to transgene-specific elements are in parentheses.

### Human stem cells (Martin Kampmann lab)

Human iPSCs in the male WTC11 background^54^ were cultured in StemFlex Medium on BioLite Cell Culture Treated Dishes (Thermo Fisher Scientific; assorted Cat. No.) coated with Growth Factor Reduced, Phenol Red-Free, LDEV-Free Matrigel Basement Membrane Matrix (Corning; Cat. No. 356231) diluted 1:100 in Knockout DMEM (GIBCO/Thermo Fisher Scientific; Cat. No. 10829-018). Routine passaging was performed as described (Tian et al. Neuron 2019). Studies with human iPSCs at UCSF were approved by the The Human Gamete, Embryo and Stem Cell Research (GESCR) Committee. Informed consent was obtained from the human subjects when the WTC11^54^ lines were originally derived.

### Ngn2-neuron Differentiation (Martin Kampmann lab)

Human iPSC-derived neurons were pre-differentiated and differentiated as described^21^. Briefly, iPSCs were pre-differentiated in Matrigel-coated plates or dishes in N2 Pre-Differentiation Medium containing the following: KnockOut DMEM/F12 as the base, 1× MEM non-essential amino acids, 1× N2 Supplement (Gibco/Thermo Fisher Scientific, cat. no. 17502-048), 10 ng ml^−1^ of NT-3 (PeproTech, cat. no. 450-03), 10 ng ml^−1^ of BDNF (PeproTech, cat. no. 450-02), 1 μg ml^−1^ of mouse laminin (Thermo Fisher Scientific, cat. no. 23017-015), 10 nM ROCK inhibitor and 2 μg ml^−1^ of doxycycline to induce expression of mNGN2. After 3 d, on the day referred to hereafter as Day 0, pre-differentiated cells were re-plated into BioCoat poly-D-lysine-coated plates or dishes (Corning, assorted cat. no.) in regular neuronal medium containing the following: half DMEM/F12 (Gibco/Thermo Fisher Scientific, cat. no. 11320-033) and half neurobasal-A (Gibco/Thermo Fisher Scientific, cat. no. 10888-022) as the base, 1× MEM non-essential amino acids, 0.5× GlutaMAX Supplement (Gibco/Thermo Fisher Scientific, cat. no. 35050-061), 0.5× N2 Supplement, 0.5× B27 Supplement (Gibco/Thermo Fisher Scientific, cat. no. 17504-044), 10 ng ml^−1^ of NT-3, 10 ng ml^−1^ of BDNF and 1 μg m^−1^ of mouse laminin. Neuronal medium was half-replaced every week. Full protocols are available on protocols.io (dx.doi.org/10.17504/protocols.io.bcrjiv4n).

At each time point, cells were washed with 1mL DPBS and then 200uL accutase was added for 7 minutes. Cells were resuspended in DPBS+rock inhibitor, and strained through a 70um filter and subjected to flow cytometry.

### 6TF-Microglia Differentiation (Martin Kampmann lab)

Microglia were differentiated as in the previous paper^26^. Briefly, iPS cells containing six doxycycline-inducible transcription factors: CEPBA, CEBPB, IRF5, IRF8, MAFB, PU.1 were passaged as a single cell suspension using accutase (Thermo Fisher Scientific A11105-01) onto Matrigel and PDL coated plates. Cells were maintained for 2 days in Essential 8 stem cell maintenance media supplemented with doxycycline and 10 uM Y-27632 (Tocris 1254). On day 2 post passage, media was replaced with Advanced DMEM/F12, 1x Glutamax, 2 ug/mL doxycycline, 100 ng/mL IL-34 (Peprotech 200-34), and 10 ng/mL GM-CSF (Peprotech 300-03). On day 4, media was replaced with Advanced DMEM/F12, 1x Glutamax, 2 ug/mL doxycycline, 100 ng/mL IL-34, 10 ng/mL GM-CSF, 50 ng/mL M-CSF (Peprotech 300-25), and 50 ng/mL TGFB (Peprotech100-21C). Media was replaced on d8, d12, and d15.

On each collection day (d0, d2, d4, d8, d12, d15), microglia were lifted for 10 min at 37C with TrypLE.

### Stem cell culture (Ernest Arenas lab)

Kolf2.1J human induced pluripotent stem cells and HS980 human embryonic stem cells were maintained undifferentiated in E8Flex media on laminin 521 (Biolamina)-coated plates, 20,000 cells/cm^2^ and passaged with ReLeSr (StemCell Technologies) or TryplLE select (Thermo Fisher), respectively. Kolf2.1J and HS980 stem cells were transfected with the TK4 PiggyBac vector and transposase plasmids at a ratio of 3:1 using nucleofection P3 primary cell kit (Amaxa 4D-nucleofector Lonza), 3 μg total DNA to 800,000 cells. Transfected cells were selected using 2 mg/ml Puromycin and stable polyclonal batches were cryopreserved.

### Midbrain Dopaminergic Neuron Differentiation (Ernest Arenas lab)

Kolf2.1J and HS980 stem cells were seeded at a density of 200,000 cells/cm2 on Geltrex-(Life Technologies) and Laminin 511-(Biolamina) coated wells and differentiated towards midbrain dopaminergic neurons according to the Arenas Lab protocol in the previous paper^27^. At day 11 and day 16 cells were replated at a density of 500,000 cells/cm2 on 1 mg/mL laminin 511 coated wells (day 11) plus 15 mg/mL poly-L-ornithine (Sigma Aldrich) (day 16).

For Kolf2.1J and HS980 cell lines, parental (wild-type) and TK4 PiggyBac-expressing stem cells were maintained undifferentiated or differentiated to dopaminergic neurons and processed in parallel for FACS analysis of BFP and EGFP at days 0, 7, 14, 21 and 28. Undifferentiated stem cells were maintained in the presence of puromycin selection, whilst differentiating cells were not. Cells were dissociated into single cell suspensions using TryplE-Select (Thermo Fisher), collected in direct trypsin inhibitor (Thermo Fisher), centrifuged at 300 x g for 5 mins and resuspended at 1,000,000 cells/ ml in FACS buffer (0.5% BSA in PBS). Cells were sorted using a 100 □m nozzle with gating to define single cells (BD Facs Aria II with BD FACSDiva 9.0.1 software). BFP+ and EGFP+ cells were defined as singlet cells with signal above the negative control for each cell line (parental, non-expressing WT stem cells).

### iPSC Culture (Deborah Kronenberg-Versteeg lab)

Human induced pluripotent stem cells (iPSC) lines BIONi010-C, (European Bank for Induced Pluripotent Stem Cells, UK), and KOLF2.1J (Jackson Laboratories, USA) were maintained on Geltrex (ThermoFisher Scientific)-coated 6-well plates (Corning) in mTeSR Plus (mTeSR+, Stemcell Technologies) media until reaching ∼80 % confluency and split using ReLeSR (Stemcell Technology) during maintenance and prior further processing. For the first 24 h after splitting, 10 µM Rock Inhibitor Y27632 (Stemcell Technologies) was added to the medium to enhance cell survival. iPSCs were cultured at 37 °C, 5 % CO2 and 100 % humidity with media changes 24 h after splitting and subsequently every other day.

### Embryoid body-based Microglia Differentiation (Deborah Kronenberg-Versteeg lab)

iPSC-derived microglia (iMic) were generated following Haenseler et al.’s protocol^30^ with small modifications. In brief, iPSCs were washed once with PBS (Gibco) and detached using ReLSR for 5-6 minutes. Cells were collected in Wash Buffer (Dulbecco’s Modified Eagle’s Medium (DMEM)-F12 (Gibco) + 0.1 % BSA Fraction V (Gibco)) and the cell suspension was centrifuged at 300 g (Heraeus Multifuge 3-SR). The supernatant was aspirated, the cells were resuspended to single-cell suspension in Wash Buffer and the cell number was determined using a hemocytometer (Neubauer Zählkammer Improved, Bard).

To generate embryoid bodies (EB) of approximately the same size, 2.5 mil iPSCs per well in 2 ml EB Induction Medium (mTeSR+ + 20 ng/ml SCF (R & D Systems) + 50 ng/ml BMP4 (Miltenyi) + 50 ng/ml VEGF (Miltenyi), supplemented with 10 μM Y27632 (Stem Cell Technologies) for the first 24 hours) were seeded into an AggreWell800 plate of the 24-well format (Stem Cell Technologies), that was prepared with anti-adherence rinsing solution (Stemcell Technologies). The cells were cultivated in the microwell plate to allow the formation of EBs at 37 °C and 5 % CO2 for 5 days with partial (75 %) media changes every day.

After 5 days, the EBs were harvested and equally distributed into 12 wells of 6-well plates in 2 ml of EB Differentiation Media (X-Vivo 15 (Lonza), 2 mM Glutamax (Gibco), 0.55 mM β-mercaptoethanol (Gibco), 100 U/ml/100 μg/ml Penicillin/Streptomycin (ThermoFisher Scientific), 25 ng/ml IL-3 (Miltenyi Biotec), 100 ng/ml M-CSF (Miltenyi Biotec)) per well.

The EBs were kept in EB Differentiation Media at 37 °C and 5 % CO2 with media changes every 7 days.

After 2-3 weeks of cultivation, non-adherent microglial precursor cells (pre-iMic) started emerging and were harvested from EB cultures by collecting the supernatant cell culture medium.

### Monolayer-based Microglia Differentiation (Deborah Kronenberg-Versteeg lab)

Additionally, iPSCs were differentiated to microglia using a monolayer-based protocol established by Takata et al.^31^. In brief, iPSCs were plated in clumps at 7,000 cells/cm2 onto GTX-coated 12-well plates (Corning) in mTeSR+ supplemented with 10 µM Rock-Inhibitor. After 24 hours, medium was changed to Supplemented Stempro-34 SFM (Gibco) medium (+ 2 mM Glutamax (Gibco), 5 µM Ascorbic Acid (Sigma-Aldrich), 150 µg/ml Transferrin (Roche), 0.45 mM Monothioglycerol (Sigma-Aldrich), 100U/ml/100μg/ml Penicillin/ Streptomycin (Gibco)) for the remainder of cultivation period, with medium changes every 2 days. On Day 0, medium was supplemented with 5 ng/ml BMP4 (Miltenyi), 50 ng/ml VEGF (Peptrotech) and 2 µM CHIR99021 (Miltenyi) and cells were transferred to hypoxia conditions for the next 8 days (37 °C, 5 % CO2, 5 % O2). On Day 2, medium was supplemented with 5 ng/ml BMP4, 50 ng/ml VEGF and 20 ng/ml bFGF (Miltenyi). On Day 4, medium contained 15 ng/ml VEGF, 5 ng/ml bFGF. From Day 6 onwards, floating precursor cells started to emerge, which were collected with the supernatant, centrifuged at 300 g for 5 min and then re-administered to the culture wells. Media composition for Day 6-10: Supplemented Stempro-34 + 10 ng/ml VEGF, 10 ng/ml bFGF, 10 ng/ml IL6 (Miltenyi), 20 ng/ml IL3 (Miltenyi), 30 ng/ml DKK1 (Miltenyi), 50 ng/ml SCF (R&D Systems). On Day 8 cells were transferred to normoxic incubator conditions (37 °C, 5 % CO2, atmospheric O2). For Day 12 and 14, medium was supplemented with 10 ng/ml bFGF, 10 ng/ml IL6, 20 ng/ml IL3 and 50 ng/ml SCF. For final differentiation, the medium was supplemented with 50 ng/ml CSF1 (Miltenyi) on Days 16-20, before pre-iMics were harvested and differentiated as described below.

### Final Microglia Differentiation and Maturation (Deborah Kronenberg-Versteeg lab)

For both protocols, pre-iMics were harvested from the supernatant cell culture medium by collecting the medium and centrifuging at 300 g for 5 minutes. The supernatant was aspirated, and cells were counted using a hemocytometer.

Pre-iMics were seeded at a density of 20,000 cells/cm2 on TC-treated cell culture plates (12-well, Corning) and differentiated to microglia in microglia differentiation medium (50 % Neurobasal Medium (Gibco), 50 % DMEM-F12 (Gibco), 1x B27 Supplement with Vitamin A (Gibco), 2 mM Glutamax (Gibco), 0.1 mM β-mercaptoethanol (Gibco), 100 ng/ml IL-34 (Miltenyi Biotec), 20 ng/ml M-CSF (Miltenyi Biotec) for up to 7 days with 3 medium changes per week.

To assess, signal intensities of BFP and GFP, cells were used for FACS analysis two times per week for the duration of the differntiation process.

At each timepoint, one well of the monolayer-based differentiation protocol and ca. 5 EBs were sacrificed and analyzed in FACS.

Cells growing in monolayer (iPSCs, monolayer-based differentiation protocol, iMics) were washed once with PBS and incubated with Accutase (Gibco) for 5-6 minutes at 37 °C, washed with Wash Buffer, collected and centrifuged at 300 g for 5 min and resuspended in FACS Buffer (PBS, 0.1 % BSA Fraction V (Gibco)). EBs were collected, washed once with PBS, incubated with Accutase for 15-20 minutes at 37 °C with intermediate vortexing and afterwards processed as cells grown in monolayer. Free-floating pre-iMics were collected from the supernatant as described above and used for FACS analysis on the day of plating pre-iMics. Cells were analyzed using a Sony SH800 cell sorter.

### Stem Cell Culture (Leslie Thompson lab)

Human iPSC lines KOLF2.1J, and CS83iCTR33n1^55^ were maintained on hESC-qualified Matrigel (Corning) in mTeSR+ medium and passaged with ReLeSR (both Stemcell Technologies). For transfection, cells were dissociated with Accutase (Invitrogen) and 200,000 cells transfected with TK4 PiggyBac vector and K13-Transposase PiggyBac plasmids at a donor:transposase ratio of 3:1 (5.33μg total DNA) using Nucleofector Solution Kit 2 (Lonza) with the Nucleofector 2b device, program B-016. Cells were allowed to recover in mTeSR+ medium supplemented with CloneR (Stemcell Technologies) and transitioned to mTeSR+ containing 1μg/ml puromycin 48hr post transfection.

### Striatal Neuron Differentiation (Leslie Thompson lab)

Neuronal differentiation was performed once iPSC colonies reached 60–70% confluency as previously described^32^. Differentiation was initiated (Day 0) by washing iPSC colonies with phosphate-buffered saline pH 7.4 (PBS – Gibco) and switching to SLI medium (Advanced DMEM/F12 (1:1) supplemented with 2 mM GlutamaxTM (Gibco), 2% B27 without vitamin A (Life Technologies), 10 µM SB431542, 1 µM LDN 193189 (both Stem Cell Technologies), 1.5 µM IWR1 (Tocris)) with daily medium changes. Four days later, cells were pretreated with 50nM Chroman 1 (MedChem Express), washed with PBS, and passaged 1:2 with StemPro Accutase (Invitrogen) for 5 minutes at 37 °C. Cells were replated onto plates coated with hESC qualified matrigel (1 h at 37 °C) in SLI medium containing CEPT cocktail (50 nM Chroman 1, 5 µM Emricasan (Selleck Chemicals), Polyamine Supplement (1:1000, Sigma-Aldrich), 0.7 µM trans-ISRIB (R&D Systems)) for 1 day after plating and continued daily feeding with SLI medim. At day 8, cells were passaged 1:2 as above and replated in LIA medium (Advanced DMEM/F12 (1:1) supplemented with 2 mM GlutamaxTM, 2% B27 without vitamin A, 0.2 µM LDN 193189, 1.5 µM IWR1, 20 ng/ml Activin A (Peprotech)) with CEPT cocktail for 1 day after plating and daily feeding was continued through day 16. At day 16, cells were pretreated with 50nM Chroman 1, dissociated with Accutase, and plated in 6-well Nunclon plates coated with PDL and hESC Matrigel at 1 × 10^6^ per well in SCM1 medium (Advanced DMEM/F12 (1:1) supplemented with 2 mM GlutamaxTM, 2% B27 (Invitrogen), 10 µM DAPT, 10 µM Forskolin, 300 µM GABA, 3 µM CHIR 99021, 2 µM PD 0332991 (all Tocris), 1.8 mM CaCl2, 200 µM ascorbic acid (Sigma-Aldrich), 10 ng/ml BDNF (Peprotech)). Medium was 50% changed every 2-3 days. On day 23, medium was fully changed to SCM2 medium (Advanced DMEM/F12 (1:1): Neurobasal A (Gibco) (50:50) supplemented with 2 mM GlutamaxTM), 2% B27, 1.8 mM CaCl2, 3 µM CHIR 99021, 2 µM PD 0332991, 200 µM ascorbic acid, 10 ng/ml BDNF) and 50% medium changes performed every 2–3 days. Cells were considered mature at day 37.

### Hypothalamic neuron differentiation (Florian Merkle lab)

KOLF2.1J hiPSCs were maintained in StemFlex iPSC medium (Thermo Fisher Scientific) on 6-well plates coated with Geltrex™ (Thermo Fisher Scientific). Cells were kept in humidified incubators at 37°C and 5% CO2, fed daily with full media changes, and passaged at 60-70% confluence with 0.5 mM EDTA for 5 min at 37°C. Upon removal of EDTA, cells were resuspended and re-plated in StemFlex medium containing 10 µM Y-27632 onto Geltrex™ - coated plates at split ratios of 1:10 – 1:12. KOLF2.1J hiPSCs were dissociated into single cell suspension using StemPro™ Accutase™ Cell Dissociation Reagent (Thermo Fisher Scientific) for 5 min at 37°C. Cells were then washed and collected with StemFlex + 10 µM Y-27632 (DNSK International) before being centrifuged at 300xg for 4 mins. The supernatant was removed, and the cell pellet was resuspended in StemFlex + 10 µM Y-27632. A cell count was then performed. KOLF2.1J hiPSCs were then seeded at 100k/cm^2^ in StemFlex + 10 µM Y-27632 in 12-well plates on Day -1. Differentiation was carried out according as previously described (Chen, Mazzaferro, et al. 2023). On Day 0, the medium was changed to the differentiation medium of N2B27 containing 100 nM LDN-193189 (DNSK International), 10 µM SB431542 (DNSK International), and 2 µM XAV939 (DNSK International). Medium was changed every other day as described in the tables below until Day 13, to generate hypothalamic neural progenitor cells (NPCs). At Day 13, NPCs were dissociated into single cell suspension using StemPro™ Accutase™ Cell Dissociation Reagent and seeded in N2B27 medium + 10uM Y-27632 into 12-well plates at 3×10^5^ cells per cm^2^. On Day 14, the medium was changed to Synaptojuice 1 (SJ1) maturation medium supplemented with 10 ng/ml Brain-derived Neurotrophic Factor, BDNF (PeproTech). The cells were maintained in SJ1 + 10 ng/ml BDNF for a week with medium changes every other day. On Day 21, the medium was switched to Synaptojuice 2 (SJ2) medium with 10 ng/ml BDNF and maintained in this medium with media changes every second day until Day 30 to generate functional hypothalamic neurons. At different time points along the differentiation timeline (Day -1 to Day 28), cells were dissociated into single cell suspensions in FACS buffer made up of DPBS (-/-) (Thermo Fisher Scientific) with 0.2% BSA (Sigma-Aldrich), 1 mM EDTA (Thermo Fisher Scientific) and 25 mM HEPES (Thermo Fisher Scientific) before being run through a BD Fortessa Cell Analyser. 3 technical replicates per sample, containing 1 million cells each were run in the cell analyser at each time point. 10,000 events were captured per technical replicate. In the FSC/SSC plot, cells were selected using a gate high for FSC and SSC to exclude debris. Separate histogram plots of BFP (violet laser 450/50) and GFP (blue laser 530/30) were generated from the gated cell population to show the cells positive for each fluorophore.

### Cortical neuron differentiation (Birgitt Schüle lab)

Human induced pluripotent stem cell (iPSC) reference line KOLF2.1J and line SUSL25 carrying a LRRK2 G2019S mutation associated with Parkinson’s disease were cultured in StemFlex media (Thermo Fisher Scientific, A3349401) on Matrigel substrate (Thermo Fisher Scientific, CB-40230) diluted in KnockOut DMEM (1:90, Thermo Fisher Scientific, 10829018). Cells were routinely passaged with ReLeSR (StemCell Technologies, 05872) at 80% confluence with supplementation of media with 1 uM of ROCK inhibitor Thiazovivin (ReproCell, 04-0017). Prior to transfection and differentiation, both iPSC lines were passaged at least twice under these conditions and synchronized to be transfected on the same day. Both iPSC lines were confirmed to be healthy and free of spontaneous differentiation as evidenced by microscopy observations of tightly packed colonies with distinctive edges, high nucleus-to-cytoplasm ratios, and prominent nucleoli.

On the day of transfection, both iPSC lines were dissociated into single cells with Accutase (Fisher Scientific, NC9464543), counted, and plated in a Matrigel-coated 6-well plate at a cell density 150,000 cells/cm2 in 1.5 ml of mTesR1 (STEMCELL Technologies, 85850) media supplemented with 1X CEPT (Chroman 1 (MedChem Express, HY-15392); Emricasan (Selleck Chemicals, S7775); Polyamine Supplement (Sigma-Aldrich, P8483-5ML); Trans-ISRIB (Tocris, 5284)). The use of CEPT in cell culture was previously described^56^. Cells were incubated for 2 hours at 37°C with 5% CO2. After 2 hours, the cells were washed with PBS without Ca2+ and Mg2+ (Fisher Scientific, 70-011-044) and the media was changed to fresh 1.5 ml of mTesR1 supplemented with 1X CEPT. The transfection mix was added dropwise to both cell lines and evenly distributed in the media by swirling the plate and performing quick side-to-side shaking of the plate. Cells were incubated for 24 hours at 37°C with 5% CO2.

After 24 hours post-transfection, the media was switched to 2 ml StemFlex with 1X CEPT. After 48 hours post-transfection, the media was changed to 2 ml StemFlex to allow cells to recover. After 72 hours post-transfection, the media was changed to 2 ml StemFlex supplemented with 2 ug/ml Puromycin (Invivogen, ant-pr-1). Cells were cultured in StemFlex with Puromycin for at least two passages before differentiation, excluding the addition of Puromycin on the day of passaging and supplementing media with 1X CEPT. After confirming stable expression of GFP by FACS, cells were taken off Puromycin and maintained in StemFlex.

For differentiation of iPSCs to neuronal progenitor cells (NPCs) on day -1, both iPSC lines were dissociated into single cells with Accutase, counted, and plated in a Matrigel-coated 6-well plate at a cell density 350,000 cells/cm2 in 2 ml of media mix that consisted of 50% StemFlex and 50% Neuronal Maintenance Media (NMM) base (Neurobasal™ Medium (serves as base, Thermo Fisher Scientific, 21103049); DMEM/F12, GlutaMAX Supplement (serves as base, Thermo Fisher Scientific, 10-565-018); Beta-Mercaptoethanol (0.1 mM, Thermo Fisher Scientific, 21985023); Penicillin-Streptomycin (1X, Thermo Fisher Scientific, 15140-122); NEAA (1X, Thermo Fisher Scientific, 11140050); GlutaMAX (1X, Thermo Fisher Scientific, 35050-061); Insulin (0.22 uM, added after filtering, Sigma-Aldrich, I9278); N2 supplement (0.5X, added fresh in weekly aliquots, Thermo Fisher Scientific, 17502-048); B27 without Vitamin A (0.5X, added fresh in weekly aliquots, Thermo Fisher Scientific, 12587010)) without the addition of insulin, N2, and B27 but with supplementation of 1X CEPT. Cells were incubated for 24 hours at 37°C with 5% CO2. After 24 hours of incubation on day 0, the media in both cell lines was changed to 4 ml/well of complete NMM (with insulin, N2, and B27) supplemented with 500 nM LDN (Peprotech, 1066208) and 10 uM SB 431542 (Peprotech, 3014193). Cells were incubated for 48 hours at 37°C with 5% CO2. Media changes to 4 ml/well of NMM with LDN and SB 431542 occurred every other day on days 2, 4, and 6. On day 7, media was changed to 3 ml/well of NMM supplemented with LDN and SB 431542. The first week of differentiation of iPSCs to cortical neurons is a sensitive period when some cell lines may experience cell death and complete detachment due to starvation because of their proliferation properties. Therefore, seeding density and feeding volumes and schedule may need to be adjusted, e.g., decrease cell density and/or increase volume of media and frequency of feeding. On days 8, 9 and 10, media was changed daily to NMM supplemented with 500 nM LDN only. On day 11, media was changed to 50% complete NMM and 50% complete N2B27 media (Neurobasal™ Medium (serves as base, Thermo Fisher Scientific, 21103049); Penicillin-Streptomycin (1X, Thermo Fisher Scientific, 15140-122); NEAA (1X, Thermo Fisher Scientific, 11140050); GlutaMAX (1X, Thermo Fisher Scientific, 35050-061); N2 supplement (0.5X, added after filtering, Thermo Fisher Scientific, 17502-048); B27 without Vitamin A (0.5X, added after filtering, Thermo Fisher Scientific, 12587010)) supplemented with 500 nM LDN. On days 12, 13, and 14, the media was changed daily to 3 ml/well of complete N2B27 supplemented with 500 nM LDN. On days 15 and 16, the media was changed daily to 3 ml/well of complete N2B27 supplemented with 500 nM LDN and 1 uM CHIR (Peprotech, 2520691). On days 17, 18, and 19, the media was changed daily to 3 ml/well of complete N2B27 supplemented with 1 uM CHIR. On day 20, the media was changed to 3 ml/well of complete N2B27 supplemented with 1 uM CHIR and 1X CEPT.

Double-coated PDL/mouse laminin 12-well plate wells were prepared 24 hours in advance before detaching and replating NPCs for maturation into cortical neurons on day 20. Wells were coated with 500 ul/well of Poly-D-Lysine (PDL, Thermo Fisher Scientific, A3890401) and incubated at room temperature for 1 hour. After 1 hour of incubation, PDL was aspirated, and wells were thoroughly washed with sterile water 3 times. Wells were further coated overnight with 600 ul/well of mouse laminin (SouthernBiotech, 1415-01) diluted in PBS with Ca2+ and Mg2+ (Fisher Scientific, 14-040-133) to 15 ug/ml. The plate was parafilmed and stored at 4°C overnight and then incubated at 37°C for 1 hour before use.

On day 21, NPCs were dissociated into single cells with Accutase, counted, and plated in a 12-well plate coated with PDL/mouse laminin at a cell density 285,000 cells/cm2 in 1 ml of media mix that consisted of 50% complete N2B27 and 50% of Terminal Differentiation (TD) media (Neurobasal A Medium (serves as base, Thermo Fisher Scientific, 12349015); Penicillin-Streptomycin (1X, Thermo Fisher Scientific, 15140-122); NEAA (1X, Thermo Fisher Scientific, 11140050); GlutaMAX (1X, Thermo Fisher Scientific, 35050-061); B27 without Vitamin A (1X, added after filtering, Thermo Fisher Scientific, 12587010); DAPT (10 uM, added after filtering, Peprotech, 2634); L-Ascorbic Acid (0.2 mM, added after filtering, Peprotech, 5088177); db-cAMP (0.5 mM, added after filtering, Peprotech, 1698950); BDNF (10 ng/ml, added fresh before media change, Peprotech, 450-02); GDNF (10 ng/ml, added fresh before media change, Peprotech, 450-10) without the addition of BDNF and GDNF but supplemented with 1X CEPT. Cells were incubated for 24 hours at 37°C with 5% CO2. After 24 hours of incubation on day 22, a half-media change was performed by aspirating 500 ul of media and adding 500 ul of fresh and complete TD media. Half-changes of complete TD media were performed every other day on days 24, 26, 28, and 30 until mature cortical neurons were prepared for the final analysis on day 32. Cortical neurons can be further kept in culture with half-media changes of TD media twice a week, e.g., on Mondays and Fridays until ready for downstream applications.

